# Auditory Word Comprehension is Less Incremental in Isolated Words

**DOI:** 10.1101/2021.09.09.459631

**Authors:** Phoebe Gaston, Christian Brodbeck, Colin Phillips, Ellen Lau

## Abstract

Partial speech input is often understood to trigger rapid and automatic activation of successively higher-level representations of words, from sound to meaning. Here we show evidence from magnetoencephalography that this type of incremental processing is limited when words are heard in isolation as compared to continuous speech. This suggests a less unified and automatic word recognition process than is often assumed. We present evidence from isolated words that neural effects of phoneme probability, quantified by phoneme surprisal, are significantly stronger than (statistically null) effects of phoneme-by-phoneme lexical uncertainty, quantified by cohort entropy. In contrast, we find robust effects of both cohort entropy and phoneme surprisal during perception of connected speech, with a significant interaction between the contexts. This dissociation rules out models of word recognition in which phoneme surprisal and cohort entropy are common indicators of a uniform process, even though these closely related information-theoretic measures both arise from the probability distribution of wordforms consistent with the input. We propose that phoneme surprisal effects reflect automatic access of a lower level of representation of the auditory input (e.g., wordforms) while the occurrence of cohort entropy effects is task-sensitive, driven by a competition process or a higher-level representation that is engaged late (or not at all) during the processing of single words.

---

Speech recognition necessarily involves the access of multiple levels of representation in response to auditory input, from phonemes to wordforms to higher-level lexical-syntactic representations that link wordforms to meaning. While much about this process remains to be elucidated, research on spoken word recognition has reached broad consensus on several points. The contributions of a vast behavioral literature (reviewed by, e.g., Dahan & Magnuson (2006); McQueen (2007); Magnuson, Mirman, & Myers (2013); Magnuson (2016)) indicate an incremental, phoneme-by-phoneme process of winnowing down the phonological wordforms that are consistent with the unfolding auditory input (e.g., Grosjean (1980); Zwitserlood (1989); Allopenna et al. (1998); and following). Conceptual information associated with those wordforms can be incrementally activated (e.g., Zwitserlood (1989); Yee & Sedivy (2006); and following), and syntactic information is rapidly invoked (e.g., Marslen-Wilson & Tyler (1980); McAllister (1988); and following). This process is highly sensitive to distributional statistics, captured by word frequency (e.g., Connine et al. (1990); Dahan et al. (2001)).

The evidence leading to this consensus comes from a broad array of experimental approaches that vary in which aspects of word recognition they can most effectively probe. These approaches use stimuli that vary from sublexical phoneme sequences to natural, connected speech. Combining evidence from these different paradigms is usually guided by an assumption that there is a uniform, automatic progression of processing triggered by speech input, such that we can expect datapoints from different points in that progression to cohere. Under this assumption, simpler or single-word paradigms will straightforwardly capture the fundamental word recognition sequence in isolation, while presenting more complex input allows us to investigate how contextual information influences, for example, the speed of processing or the set of lexical candidates under consideration.

In **Figure 1**, we sketch a representative sequence of processing proposed to occur in response to each phoneme of speech input. TRACE (McClelland & Elman, 1986) is an example of a model that is consistent with the illustrated principles. Each level of representation automatically determines the most likely interpretation of the input through local competition and broadcasts this interpretation through feed-forward and feed-back connections. An assumption of automaticity implies that any speech input engages this processing hierarchy in the same manner. The task context might change the information available at different levels, but not the basic sequence of processing. However, if that assumption of automaticity is incorrect, then the basic process of word recognition could deviate significantly according to the demands of different comprehension scenarios. This deviation could occur because of variation in, for instance, the relevance of different types of information to different experimental tasks, the ease of word segmentation, or the degree to which word-to-word dependencies occur in the input.

**Figure 1.**
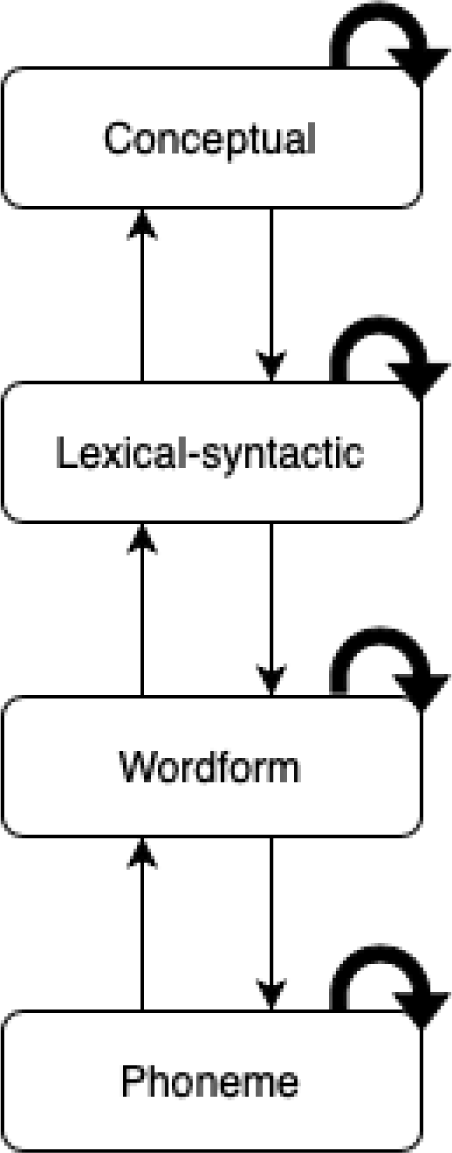
Automatic Sequence of Processing Assumed to Occur in Response to Each Phoneme of Speech Input Note. Straight arrows indicate connections between levels of representation. Curved arrows indicate a within-level competition/selection process.

In this paper we present neural evidence that word recognition in isolation may proceed in a qualitatively different way from word recognition in continuous speech. Behavioral measures or paradigms requiring an explicit response to each stimulus make comparison between isolated words and continuous speech difficult, with single trials generally measuring the status of just a single item in the lexicon. Instead, we turn to a neural measure – multivariate temporal response function (mTRF) analysis of magnetoencephalography (MEG) responses – that can be applied in exactly the same way to single-word and continuous-speech listening, and that reflects distributional properties of the entire class of word candidates consistent with each presented phoneme. We show that the effects of two measures that have both been understood to reflect automatic wordform-level processing in fact dissociate robustly according to the nature of the experiment. This dissociation implicates a break in the automaticity of the sequence of activation and indicates a difference between the processing of words presented in isolation and words presented in continuous speech. Our findings have implications for the architecture of word recognition models as well as for experimental approaches to studying speech perception.

## Phoneme Surprisal and Cohort Entropy

The neural response to speech has been shown to be modulated by information-theoretic properties of the set of wordforms that match the auditory input at any given phoneme (Brodbeck et al., 2018, 2022b; Di Liberto et al., 2019; Donhauser & Baillet, 2020; Ettinger et al., 2014; Gagnepain et al., 2012; Gaston & Marantz, 2018; Gillis et al., 2021; Gwilliams et al., 2018, 2020; Gwilliams & Marantz, 2015; Kocagoncu et al., 2017). Two of these properties in particular – cohort entropy and phoneme surprisal – have emerged as promising means of investigating the time course of auditory word recognition.

*Phoneme surprisal* at a given phoneme is a measure of how much information that phoneme provides for identifying the current word. It is defined as the conditional probability of that phoneme given the preceding sequence of phonemes in the current word. Phoneme surprisal at position *i* in a wordform is defined as −*log*_2_ *p*(*k*_*i*_ | *k*_1_, … *k*_*i*−1_) where *k*_*i*_ is the phoneme at position *i* and *i* = 1 for the first phoneme in the wordform. *Cohort entropy* at that same phoneme, in contrast, is a measure of how much uncertainty there is across word forms that match the phoneme sequence up to the current phoneme. It is determined by the probability distribution over wordforms that might complete that phoneme sequence. Cohort entropy at position *i* in a wordform is defined as 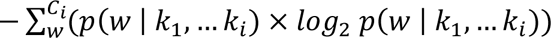 where *w* is each wordform in the cohort *Ci* of wordforms consistent with the sequence of phonemes *k*_1_, … *k*_*i*_. One of the critical differences between these formulations is that cohort entropy is forward-looking in a way that phoneme surprisal is not. A cohort entropy effect reflects expectations for potential wordform candidates that would be consistent with the current input, while a phoneme surprisal effect may only reflect the degree to which previously formed representations are updated based on the new input (see Pickering and Gambi (2018) on entropy effects as strong evidence for prediction).

More neural activity is generally observed in response to higher surprisal, or lower probability, phonemes, consistent with many cognitive domains in which predictable or higher probability stimuli elicit reduced neural responses (see Aitchison & Lengyel (2017)). Exactly how cohort entropy should be expected to drive neural activity in this case is less clear, though there is evidence for the relevance of the broader concept of entropy across a range of areas within cognitive neuroscience (Bestmann et al., 2008; Crupi et al., 2018; Friston, 2005; Hale, 2016; Strange et al., 2005; Weissbart et al., 2020; Whiteley & Sahani, 2008; Willems et al., 2016). A larger set of word candidates has a higher cohort entropy than a smaller set of candidates, but the size of the candidate set is not the only determinant of uncertainty; a set of candidates in which probability is equally distributed has a higher cohort entropy than a set of candidates in which probability is concentrated on a single candidate. Greater uncertainty among word candidates could be associated with more neural activity due to an intensified process of lexical competition (Gagnepain et al., 2012), or due to increased attentional gain on bottom-up input (Donhauser & Baillet, 2020), or it could be instead that lower uncertainty is a precondition for other processes to be engaged (Ettinger et al., 2014).

Despite these differences, phoneme surprisal and cohort entropy are often investigated and presented in tandem as interchangeable indicators of wordform-level processing. One likely reason for this approach in the literature is that the conditional probabilities underlying both measures are calculated from the same probability distributions of wordforms consistent with the input. The two variables are also often correlated, and their effects in neural data frequently co-occur. Finally, in a hypothesized model of word recognition that includes automatic engagement of successive representational levels regardless of task or context, phoneme surprisal and cohort entropy effects are simply two different windows into the same automatic flow of activation through the system.

## Variation in Neural Effects of Cohort Entropy and Phoneme Surprisal

Despite frequently being treated interchangeably, a careful look at the prior literature reveals considerable variation in whether and when phoneme surprisal and cohort entropy effects manifest across experiments. This variation has not previously been examined systematically.

Thus, before we proceed to our own study, we review this literature and consider whether there are properties of the stimulus or experimental context that can help explain when cohort entropy and phoneme surprisal effects do or do not occur, and what this might mean for the processes and levels of representation they describe. An account of this variability is important for improving the utility of phoneme surprisal and cohort entropy as measures for investigating speech perception and specifically the class of active items in competition for recognition at any given point in a word. However, understanding this variability also has the potential to illuminate dissociable sub-processes in word recognition.

We begin by trying to characterize why cohort entropy and phoneme surprisal effects occur at all in some experiments and not in others, though further efforts to understand variation in the localization and time course of these effects will also be important. In **Table 1**, we summarize existing electrophysiology (primarily MEG) studies that have tested for effects of cohort entropy and phoneme surprisal on neural activity. Effects of both cohort entropy and phoneme surprisal have been reported in behavioral measures of auditory word recognition (Baayen et al., 2007; Balling & Baayen, 2012; Bien et al., 2011; Kemps et al., 2005; Wurm et al., 2006). However, testing for such effects in behavioral data generally requires constructing a cumulative measure of a phoneme-level variable across the course of the word or selecting the variable’s value at just one phoneme position as the predictor. Therefore we restrict our focus here to neural measures that have the temporal resolution to examine cohort entropy and phoneme surprisal effects on a phoneme-by-phoneme basis. We exclude two additional studies (Di Liberto et al., 2019; Gwilliams et al., 2018) which did not report effects of cohort entropy and phoneme surprisal separately, as well as a third study (Wang et al., 2021) in which results are described as being inconsistent with a cohort entropy effect, even though cohort entropy values are not used for comparison between critical conditions.

**Table 1:** *Properties of the Stimulus and Experimental Task for Existing Electrophysiology Studies Reporting Phoneme Surprisal or Cohort Entropy Effects*

| Study | Phoneme surprisal effect? | Cohort entropy effect? | Stimulus | Experimental task | Multimorphemic words included? |
| --- | --- | --- | --- | --- | --- |
| Gagnepain et al. (2012) | yes | no | single words | pause detection | no |
| Ettinger et al. (2014) | yes | yes<sup>^</sup> | single words | lexical decision | yes |
| Brennan et al. (2014) |  | no | single words | lexical decision | no |
| Lewis & Poeppel (2014) |  | no | single words | lexical decision | no |
| Gwilliams & Marantz (2015) | yes |  | single words | lexical decision | yes |
| Kocagoncu et al. (2017) |  | yes<sup>^</sup> | single words | nonword detection | not specified |
| Gaston & Marantz (2018) | yes | yes<sup>*</sup> | three-word phrases | phrase acceptability | yes |
| Brodbeck et al. (2018) | yes | yes | continuous speech | comprehension questions | yes |
| Donhauser & Baillet (2020) | yes |  | continuous speech | comprehension questions | yes |
| Gwilliams et al. (2020) | yes | yes | continuous speech | comprehension questions | yes |
| Gillis et al. (2021) | yes | yes | continuous speech | comprehension questions | yes |
| Brodbeck et al. (2022b) | yes | yes | continuous speech | comprehension questions | yes |

*Note.* Grey cells indicate studies that did not test for the specified effect. Superscripts indicate that a reported cohort entropy effect did not survive when phoneme surprisal was controlled for (\*), or that such a test was not performed (^).

**Table 1** demonstrates that phoneme surprisal and cohort entropy effects have very different profiles across studies. Phoneme surprisal effects were reported in all studies that tested for them (Brodbeck et al., 2018, 2022b; Donhauser & Baillet, 2020; Ettinger et al., 2014; Gagnepain et al., 2012; Gaston & Marantz, 2018; Gillis et al., 2021; Gwilliams et al., 2020; Gwilliams & Marantz, 2015), and thus appear to be robust to variation in stimulus and experimental task. Cohort entropy, in contrast, produces mixed results. Among studies that presented single words and short phrases, three reported cohort entropy effects (Ettinger et al., 2014; Gaston & Marantz, 2018; Kocagoncu et al., 2017) and three tested for but failed to find them (Brennan et al., 2014; Gagnepain et al., 2012; Lewis & Poeppel, 2014). The presence or absence of multimorphemic words (words comprised of a root and at least one affix) in the study is potentially relevant, as the three studies that failed to find cohort entropy effects included only monomorphemic words. However, more important in our view is that the three single-word studies that reported cohort entropy effects did not exclude the possibility that these effects were due to the highly correlated phoneme surprisal measure. Gaston and Marantz (2018) in fact found that their significant cohort entropy effect was no longer significant in a model that controlled for phoneme surprisal, and the other two studies (Ettinger et al., 2014; Kocagoncu et al., 2017) did not conduct such a test. In continuous speech, cohort entropy effects were reported in all studies that tested for them (Brodbeck et al., 2018, 2022b; Gillis et al., 2021; Gwilliams et al., 2020), with methods that controlled for effects of phoneme surprisal. We conclude that, in the existing electrophysiology literature on speech recognition, there is strong evidence for phoneme surprisal effects across the board, but for cohort entropy effects only in continuous speech.

A true dissociation between cohort entropy and phoneme surprisal effects would indicate not only that these measures index different levels of representation or processes, but also that the activity that drives cohort entropy effects may be reduced or occur not at all during the processing of single words, or at least not incrementally (i.e., phoneme by phoneme). This is not consistent with all processing steps being engaged in a fully automatic sequence during speech recognition. However, this interpretation of the prior literature is complicated by the fact that many of these studies did not control for potential confounds, such as acoustic variables and overlapping responses to different phonemes. Differences in statistical power or analysis methods (which vary widely) may also have contributed to the apparent influence of stimulus on cohort entropy effects.

## The Current Study

Hypothesizing that cohort entropy and phoneme surprisal do, indeed, dissociate, and that cohort entropy effects do not occur for single words, we evaluated cohort entropy and phoneme surprisal effects on the neural response to speech in a simple single-word paradigm and then directly compared this data to an existing continuous-speech dataset (Brodbeck et al., 2022b).

Comparing single-word and continuous-speech data requires that the two types of responses be evaluated with the same method. Analysis techniques traditionally applied to single-word studies are not suitable for responses to continuous speech, and generally fail to account for acoustic and other confounding variables, as well as the overlapping nature of phoneme responses. Instead, we modeled source-localized MEG data with multivariate temporal response functions (**Figure 2**), a method that deals with acoustic confounds and was originally developed for continuous speech. This allowed for novel comparison between single words and continuous speech as well as a more accurate characterization of the single-word response relative to previous analyses.

**Figure 2.**
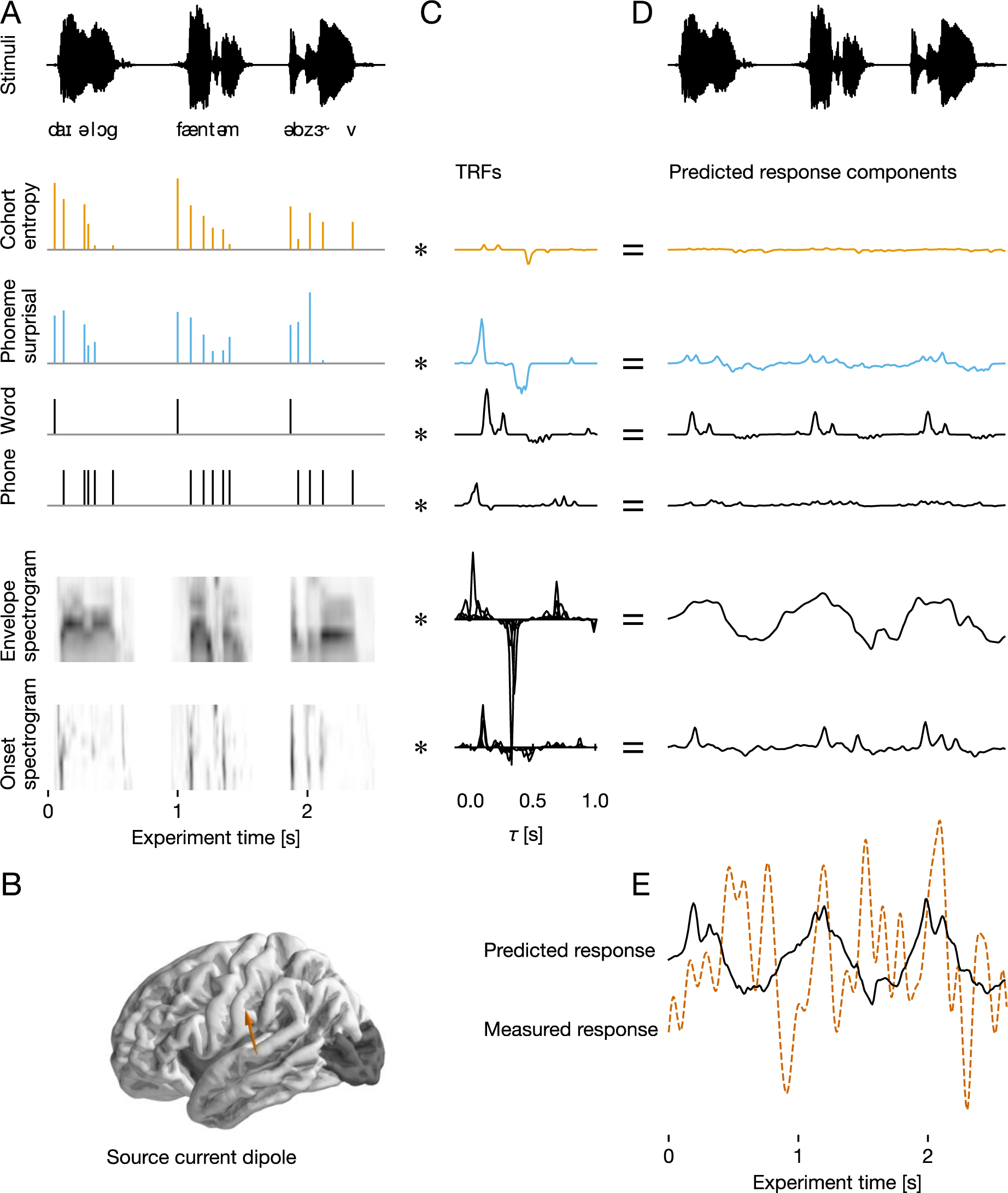
Multivariate Temporal Response Function (mTRF) Analysis of Magnetoencephalography Data Note. Multivariate temporal response function (mTRF) analysis models brain activity as a continuous response to multiple concurrent predictor variables that describe the sequence of words. mTRF models were estimated separately for each subject, and the resulting model fits were analyzed at the group level. **(A)** For mTRF analysis, brain responses were analyzed as continuous recordings, temporally aligned with the stimulus sequence of words, presented with a 267 ms inter-stimulus interval. Predictor variables that quantify different properties of the stimuli were also continuous time series, aligned with the stimuli. Cohort entropy and phoneme surprisal were generated using impulses at phoneme onsets, scaled by the relevant quantity. Covariates included word and phoneme onsets, an 8-band auditory envelope spectrogram (i.e., eight predictors reflecting different frequency bands), and an 8-band auditory onset spectrogram. **(B)** Neural activity was quantified as distributed minimum norm current estimates, i.e., estimated currents at a grid of dipoles covering the cortical surface. The analysis was restricted to the temporal, frontal, and parietal lobes (the dark shading indicates regions excluded from the analysis). One dipole from one representative subject is used in this figure for illustration (brown arrow). **(C)** TRFs were estimated using a coordinate descent algorithm to predict the neural signal from the predictor variables. Each predictor (A), convolved with the corresponding TRF (B), equals to a component of the predicted response **(D)**. These response components are thus again aligned with the stimulus, repeated at the top of (D). TRFs were estimated jointly, i.e., each TRF, convolved with its corresponding predictor variable, predicted a component of the neural activity, and the sum of these component responses is the predicted brain response **(E)**. Model performance was evaluated by the proportion of the variability in the measured response that was explained by the predicted response, using 5-fold cross-validation.

Participants heard a list of 1000 monomorphemic words with an inter-stimulus interval of 267 ms, and were asked to respond to randomly occurring semantic relatedness probes to ensure attention and motivate higher-level lexical processing (see Brodbeck et al. (2018) on the lack of lexical effects in unattended speech). Models were fit using 5-fold cross-validation in each subject separately. We evaluated the models by the proportion of variability they explained in the source-localized MEG recordings, correcting for multiple comparisons using threshold-free cluster enhancement (Smith & Nichols, 2009). Unless noted specifically, analyses were performed on the surface of the temporal, frontal and parietal lobes combined (excluded areas are shaded in **Figure 2B**).

## Materials and Methods

### Participants

We collected MEG data from 24 people. Sample size was chosen in accordance with the previous studies cited in **Table 1**. All participants were right-handed, native speakers of English, and seven were also native speakers of additional languages. None reported a history of neurological or linguistic impairment, brain injury, or hearing loss. All reported normal or corrected-to-normal vision. The procedure was approved by the Institutional Review Board at the University of Maryland, College Park and all participants provided written informed consent. Participants were compensated with their choice of $15 or one course credit per hour of participation. The full session (including another, unrelated study) lasted two hours.

One dataset was excluded before data processing because of participant fatigue and an earbud falling out during the experiment. After this exclusion, we computed accuracy on the semantic relatedness task and excluded any participant with accuracy lower than a cutoff one standard deviation below the mean. This excluded three of 23 participants, and provides assurance that the included participants were accessing lexical information above the wordform level. After preprocessing, two additional datasets were excluded due to excessive magnetic noise. 18 datasets are therefore included in our analysis. Raw data is available at <u>doi:10.18112/openneuro.ds004276.v1.0.0</u>.

### Stimuli

Our stimuli were word recordings from the Massive Auditory Lexical Decision (MALD) database (Tucker et al., 2019), which includes the timing of phoneme boundaries as determined by a forced aligner. The set of 1000 words we selected had no missing variables in the database and were monomorphemic per MALD, CELEX (Baayen et al., 1995), and first author judgment. We excluded all items with the following labels in MALD: Preposition, Interjection, Name, Unclassified, Conjunction, Pronoun, Determiner, Letter, Not, Ex, Article, To. We also removed items with the 10% lowest frequency values, and excluded homophones, inappropriate and particularly evocative words, and any item for which the pronunciation in the recording was noticeably divergent from American English. The full lists of stimuli and semantic relatedness probes (see below), as well as associated stimulus variables from MALD, are available at <u>osf.io/u56ea/</u>.

### Procedure

The study was always the second of two experiments in a session. Before the MEG recording, we used a Polhemus 3SPACE FASTRAK to digitize participant head shapes as well as the positions of five affixed marker coils. These marker coils were used to record head position relative to the MEG sensors before and after each study in the session. We recorded continuous MEG data, inside a magnetically shielded room, with a 160-channel axial gradiometer whole-head system (Kanazawa Institute of Technology, Kanazawa, Japan). Our sampling rate was 1000 Hz, and we used an online 60 Hz notch filter and 200 Hz low-pass filter.

Participants lay supine and looked at a screen overhead, while holding a button box in each hand. They wore foam earbuds and volume was adjusted to their comfort level. We instructed participants that they would hear a long series of random words, and that they should simply listen to the words while watching for probe words that would randomly appear on the screen with a question mark. They were instructed to press a button (with left hand for No and right hand for Yes) to indicate whether the word on the screen was related in any way to the word they had heard just before it. Because probe words were unpredictable, good performance on the task requires lexical-syntactic and conceptual information access to have occurred for most stimuli.

We used Presentation (Neurobehavioral Systems, Inc., www.neurobs.com) to present the experiment. Our parameter and scenario files are available on OSF (https://osf.io/u56ea/). There were 1000 auditory trials interspersed pseudo-randomly with 97 semantic relatedness probe trials. The amount of time between trials was 267 ms. A visual fixation cross was on screen continuously during auditory trials and during the inter-trial interval. Each auditory trial simply consisted of presentation of the auditory stimulus and lasted the length of the auditory stimulus. Visual probe trials were pseudo-randomly distributed with a maximum interlude of 20 trials between probes. The probe (e.g., “podium?”) stayed on the screen until the participant pressed a button to answer.

We selected this task so that it would apply equally well to all types of words, and because we did not want button presses to occur on critical trials (as would happen in, e.g., lexical decision). The probe trials for which we expected participants to answer “No” were selected randomly from the list of eligible words that we did not end up using for auditory trials. Probe trials for which we expected participants to answer “Yes” were synonyms taken from the WordNet (https://wordnet.princeton.edu) page of the preceding auditory item and were also monomorphemic so as not to be trivially distinguishable from “No” trials. There was no overlap between probe words and words used in auditory trials. Which auditory trials would be followed with a probe were randomly selected. “Yes” and “No” probes were equally distributed.

The experiment lasted roughly 17 minutes. There was no built-in break, but participants were instructed that if they wished to take a break, they should simply delay their button press on a probe trial.

### Data Preprocessing

We processed the data using mne-python version 0.22 (Gramfort et al., 2013, 2014) and Eelbrain 0.34 (Brodbeck et al., 2019). Code for processing and analysis can be accessed via <u>osf.io/u56ea/</u>.

During file conversion with mne-python’s kit2fiff GUI, we excluded any faulty marker measurements. We co-registered each digitized head shape with the Freesurfer (Fischl, 2012) “fsaverage” brain, using mne-python’s co-registration GUI. We first used rotation and translation to align the digitized head shape and average MRI by the three fiducial points. We then used rotation, translation, and 3-axis scaling to minimize the distance between digitized head shape and average MRI points using the iterative closest point (ICP) algorithm. Convergence was always achieved within 40 iterations. For one participant, outlying points on the digitized head shape were removed between fitting to the fiducials and applying ICP.

Flat channels were automatically removed, and we used temporal signal space separation (Taulu & Simola, 2006) for removal of extraneous artifacts, with a buffer duration of 10 seconds. We then band-pass filtered the recordings between 1 – 40 Hz (mne-python default settings) and used ICA (independent components analysis, with extended infomax method) for removal of ocular, cardiac, and other extraneous artifacts. Components were selected manually based on their topography and time-course. After removing artifactual ICA components, we further low-pass filtered the data at 20 Hz, cropped it from 1 s before the first word to 2 s after the last word, and down-sampled it to 100 Hz.

To compute a noise covariance matrix, we used two minutes of empty room data recorded before or after each session. We defined the source space on the white matter surface with a four-fold icosahedral subdivision, with 2562 sources per hemisphere. Orientation of the source dipoles was fixed perpendicular to the white matter surface. Continuous data were source localized with the regularized minimum norm estimator (λ = 1/6). The use of signed current estimates ensures that the expected (mean) value of the noise is 0, making this method suitable for single-trial source localization.

## Analysis

### Behavioral Data

Mean accuracy was computed after the exclusion of one participant a priori. The mean number of correct probe responses was 73.6 (out of 97) with a standard deviation of 18.4. The number of correct probe responses was lower than one standard deviation below the mean for three participants, so they were excluded from further analysis. One participant answered 13 of 97 probes correctly. We kept this participant in the dataset because this was so far below chance that the only plausible explanation seemed to be that they had reversed which hand they were supposed to use to make Yes and No responses.

### Predictors for Neural Data

For each acoustic or linguistic variable of interest used as a predictor of the neural response (see list below), a time series was created indicating the value of the predictor at each time point in the experiment. Our study did not actually present a single continuous stimulus (rather, we presented individual words with short intervening pauses), but a single time series reflecting predictor values (or pauses) throughout the entire experiment could still be created (see **Figure 2A**). Probe trials were modeled simply as silence. The timing of phoneme onsets was taken from the forced aligner information made available with the MALD recordings.

For acoustic predictors (envelope and onset spectrogram), the value of the predictor could vary continuously at each time point of the stimulus. Linguistic predictors consist of impulses at phoneme onsets only, and thus have a value of zero at all other points in the stimulus. Of the lexical predictors, phoneme onset and word onset each consist of binary impulses, while entropy and phoneme surprisal consist of impulses that are scaled continuously according to the entropy or surprisal value of that phoneme.

### Acoustic Envelope Spectrogram

A gammatone spectrogram (Heeris, 2018) was computed for each stimulus waveform with 256 channels regularly spaced in ERB space between 20 and 5000 Hz. These spectrograms were resampled to 100 Hz to match the MEG data and binned into eight equally spaced frequency bands.

### Acoustic Onset Spectrograms

The high resolution gammatone spectrograms were processed with an algorithm for acoustic edge extraction (Brodbeck et al., 2020; Fishbach et al., 2001). The onset spectrograms were also resampled to 100 Hz and binned into eight bands.

### Word Onsets

Word onsets were represented as a single, equally valued impulse at the onset of every word, as determined from the forced alignments. These were included to control for responses that uniformly occur time-locked to speech onset for all words.

### Phoneme Onsets

Phoneme onsets (excluding phonemes that were also word onsets) were represented as equally valued impulses on a single predictor time series that included all remaining phoneme positions. These were included to control for responses that occur time-locked from phoneme onset but do not scale with surprisal and entropy.

### Phoneme Surprisal and Cohort Entropy

These variables were calculated based on an implementation of the cohort model of word perception (Marslen-Wilson, 1987), as in Brodbeck et al. (2018). Initially, a dictionary was created combining frequency information from SUBTLEX (Brysbaert & New, 2009) with pronunciations (phoneme sequences) from the CMU pronouncing dictionary (Weide, 1994), adding any pronunciations for stimuli that were missing from the CMU dictionary. This dictionary was then used to compute the set of words compatible with the input so far for each word at each phoneme position. These cohorts, together with the SUBTLEX frequencies, were used to compute a probability distribution over possible words for each phoneme position. The *cohort entropy* predictor contained an impulse at each phoneme onset, scaled by the entropy of that cohort. The *phoneme surprisal* predictor contained an impulse at each phoneme onset scaled by the surprisal of that phoneme, based on the posterior probability of that phoneme given the preceding phoneme’s cohort.

### mTRF Analysis

An mTRF maps a set of predictor variables to a single outcome time series. Here, independent mTRFs were estimated for each subject and for each virtual current source of source-localized MEG data (see **Figure 2**). The neural response at time *t*, represented as *y_t_*, is predicted jointly from *N* predictor time series, represented as *x_i,t_*, convolved with a corresponding mTRF, represented as *h_i,τ_*, with weights for all N predictors at a range of delays *T*:

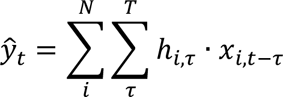

mTRFs were generated from a basis of 50 ms wide Hamming windows centered at *T_basis_*=[- 100,…,1000) ms. All responses and predictors were standardized by centering and dividing by the mean absolute value.

For a given set of predictors, the predictive power was estimated through 5-fold cross-validation. For this, the continuous data and corresponding predictors were split into five contiguous partitions of even length. The neural responses of each partition were predicted with an mTRF trained on the remaining four partitions to minimize *l*1 error. Within each set of four training partitions, each partition in turn served as validation data once. Thus four mTRFs were estimated based on coordinate descent, with early stopping based on the validation data (David et al., 2007). The validation data was used to selectively stop training predictors when they caused an increase in error in the validation set. Those four mTRFs were then averaged to predict the responses to the unseen (fifth) test segment.

For evaluating the predictive power of phoneme surprisal and cohort entropy, we compared the predictive power of the full model with that of a model that was identical except for not including the predictor under investigation. Together with the cross-validation, this assures a conservative estimate of the unique predictive power of the predictor under investigation, while controlling for the predictive power of all the other variables. The anatomical maps of explanatory power of the two models were smoothed (Gaussian kernel, SD = 5 mm) and compared using a mass-univariate related measures *t*-test based on threshold-free cluster enhancement (TFCE) (Smith & Nichols, 2009), with a null distribution based on 10,000 random permutations of condition (model) labels.

For analysis of the individual predictor TRFs, the five estimates of the TRFs from the five different test partitions were averaged in each subject. To visualize the TRF current over time, the TRF was restricted to the anatomical region of interest (ROI) defined as the area in which the surprisal predictor significantly improved predictions (*p* ≤ .05 corrected with TFCE). To visualize TRF amplitudes, the absolute values of the TRFs were averaged across the anatomical ROI (**Figure 5A**). To visualize the anatomical distribution, the absolute values of the TRF were averaged across a given time window and subjects, and the resulting images were smoothed with a Gaussian kernel (SD = 5 mm; **Figure 5B**). To visualize current direction, the TRFs were further analyzed using principal component analysis (**Figure 5C, D**). Within the same area, defined based on significance of the surprisal predictor, and separately for each hemisphere and each participant, principal component analysis was applied to the surprisal TRF, such that the TRF was decomposed into different time courses, each with a specific anatomical distribution. To visualize the dominant trend in the TRFs, the first principal component was analyzed, i.e., a single spatial topography and corresponding time course for each participant.

The advantage of this approach over the amplitude analysis is that the signed current direction can be visualized. Because the sign of a principal component is arbitrary, the components were aligned across subjects such that the average current vector was pointing upward. For components whose average current vector pointed downward, both component and time-course were multiplied by -1.

The TRF time-course was then evaluated in each hemisphere using a mass-univariate one-sample *t*-test with TFCE, with the null hypothesis that the average current direction is random (i.e., not different from 0). The null distribution was based on the maximum statistic in 10,000 random permutations of the signs. To test for hemispheric differences, a mass-univariate repeated measures *t*-test with the same parameters was used.

### Status of the First Phoneme

Two previous studies did not find evidence for surprisal and entropy effects related to the first phoneme of each word (Gaston & Marantz, 2018; Brodbeck et al., 2018). However, since neither study actually showed a significant difference between surprisal or entropy effects at first vs. subsequent phonemes, we here performed a preliminary analysis to determine whether surprisal and entropy at word-initial phonemes should be modeled separately from at subsequent phonemes. To this end, we compared the model treating all phonemes uniformly (as depicted in **Figure 2**) to a model in which surprisal and entropy at the first phoneme were modeled as separate predictors from surprisal and entropy at non-initial phonemes. The more complex model, in which they were modeled separately, was not significantly better (*t_max_* = 2.74, *p = .*341, multiple comparison correction in bilateral temporal lobe only). We therefore proceeded with the simpler model in which surprisal and entropy at initial phonemes are not modeled separately (as shown in **Figure 2**).

### Overall Model Performance

We assessed the overall performance of the full model in held-out data by averaging relevant performance metrics across subjects, and then report the maximum across the brain. The full model explained 2.9% of the variability of the source localized MEG responses at the best current dipole. A more common metric is the correlation between the predicted and the actual MEG signal, which reached *r* = 0.25, in line with previous studies.

### Comparison with Connected Speech

For this comparison, data from 12 participants listening to 47 minutes of a non-fiction audiobook were used (for more details see Brodbeck et al. (2022b); data available via Brodbeck et al. (2022a)). The two datasets had one participant in common. Data were acquired on the same MEG equipment and processed and analyzed with analogous procedures, with one exception: for estimation of the mTRF models, data were split into four partitions instead of five. This was done to speed up computations (requiring training of fewer models) and because the longer recording resulted in more training data per participant. Audiobook stimuli were labeled using the Montreal Forced Aligner (McAuliffe et al., 2017), and predictor variables were generated as for the single-word data.

The audiobook dataset included more data per participant, raising the concern that larger effect sizes would be expected just because of the larger amount of data. In total, a participant in the single-word experiment heard 1000 words with a total of 4889 phonemes, whereas the connected speech stimuli contained 27810 phonemes. To address this, we repeated the comparison between experiments with a subset of the continuous-speech data. Similar numbers of phonemes could be achieved by combining segments 5 and 6 of the audiobook stimulus (4964 phonemes) or segments 7 and 8 (4949 phonemes). 5-fold cross-validation was used for this analysis just as for the single-word experiment.

## Results

To ensure that responses reflect attentive lexical processing, we applied behavioral exclusion criteria (see Methods, though note that statistical outcomes do not change when behavioral exclusions are not applied). Subjects included in the analyses presented here responded accurately to at least 69% of relatedness probes (group mean 82.9%).

To test whether phoneme surprisal and cohort entropy improve the estimated neural response in a single-word design, we fitted three separate TRF models: the full model depicted in **Figure 2**, an (otherwise identical) model missing the surprisal predictor, and an (otherwise identical) model missing the entropy predictor. To control for responses associated with different aspects of speech processing, all models included an acoustic envelope spectrogram, an acoustic onset spectrogram, and word and phoneme onsets. We found that the full model was significantly better than the model without phoneme surprisal (*p* < .001), indicating that phoneme surprisal explains a component of the brain responses that none of the other included variables could explain. However, comparison with the model lacking cohort entropy led to no significant difference (*p* = .260, see **Figure 3A, B**). The difference between the two variables was reliable: the model improvement due to surprisal (i.e., the explanatory power of surprisal) was significantly larger than that due to entropy (*p* = .007).

**Figure 3.**
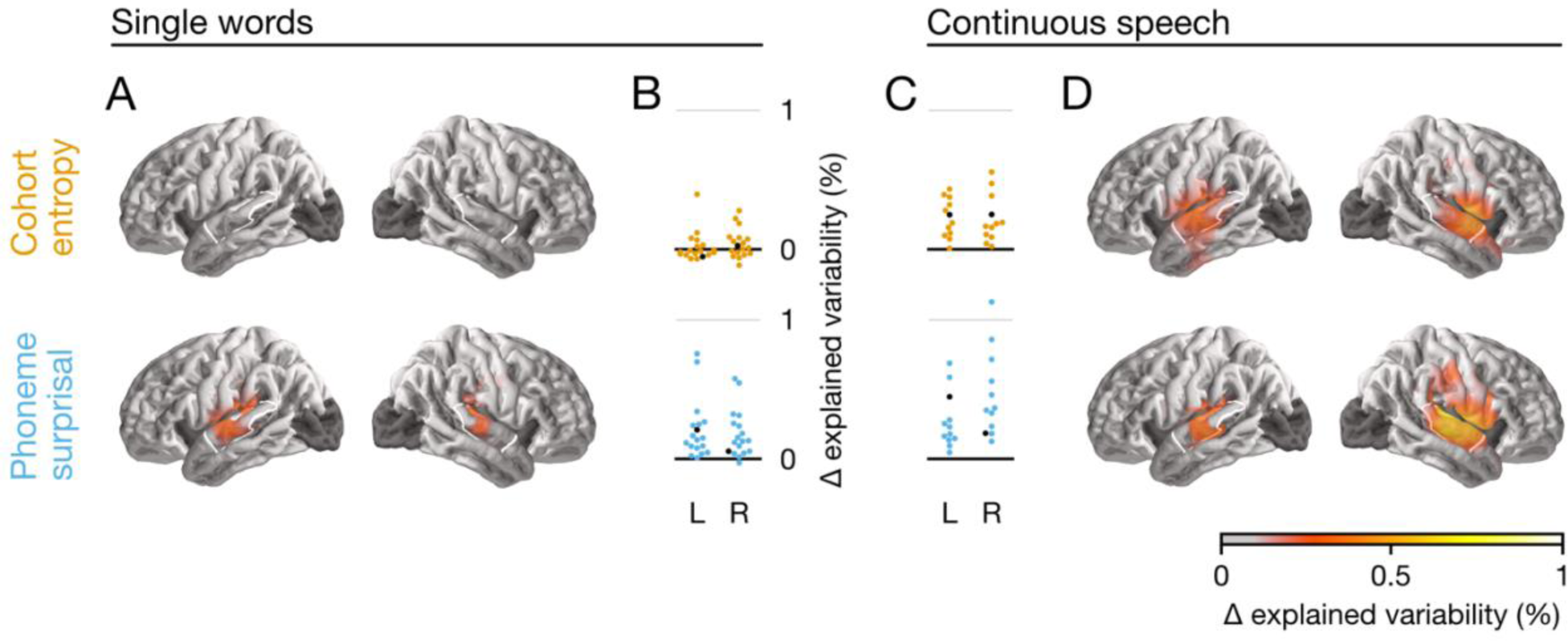
Model Evaluation and Comparison to Continuous Speech Note. The anatomical plots (**A, D**) show regions where the given predictor significantly improved the model fit (p < .05, corrected). The white outline indicates an anatomical region of interest (ROI) defined as the posterior 2/3 of the superior temporal gyrus. The swarm plots (**B, C**) show average proportion of variability in that ROI that is uniquely explained by entropy or surprisal, respectively. Each dot represents one participant. The black dot represents a single participant that took part in both experiments. While surprisal improves the model fit in both experiments in almost all participants, entropy does so only in the continuous-speech data. Explained variability (explanatory power) is expressed as percentage of the maximum variability explained by the full model in the single-word data.

This finding contrasts with previously reported results in connected speech (see **Table 1**).

To address this apparent difference, we compared our single-word data to an existing continuous-speech dataset (Brodbeck et al., 2022b) that consisted of recordings from 12 participants listening to 47 minutes of an audiobook, and had been acquired with the same MEG scanner. The correlation between the phoneme surprisal and entropy values over all phonemes was similar in the two datasets (single words: *r* = 0.39; continuous speech: *r* = 0.41). Using closely matched analysis methods, we found that, for the continuous-speech data (see **Figure 3C, D**), phoneme surprisal significantly improved the model (*p* < .001) and cohort entropy did as well (*p* < .001). In the whole brain analysis, the explanatory power of phoneme surprisal and cohort entropy did not differ significantly (*p* = .720).

To confirm that this difference between experiments was statistically meaningful, we compared the two datasets directly. We extracted the mean of the model fit metric in an anatomically defined region of interest (ROI), including the posterior two thirds of the superior temporal gyrus of each hemisphere. This value did not differ between the left and right hemisphere ROIs in any of the four categories (surprisal/entropy, single words/continuous speech; all *t* ≤ 1.74, *p* ≥ .110), so we averaged the values for the two hemispheres. The conclusion that there is a difference between experiments would follow from an interaction between cohort measure (surprisal vs. entropy) and experiment (single words vs. continuous speech). However, due to the different effect sizes between experiments, the additive model underlying ANOVA may not be appropriate. Instead, we calculated the ratio between the predictive power of entropy and surprisal for each participant, and then across participants compared this ratio between continuous speech and single words. This ratio was significantly higher for continuous speech than for single words (continuous speech *M* = 0.68, *SD* = 0.45; single words *M* = 0.10, *SD* = 0.59; *t28* = 2.80, *p* = .009). Based on this difference in ratio, we reject the null hypothesis that surprisal and entropy make equal relative contributions to the explanatory power of the models in the two experiments. Consistent with this conclusion, effect sizes for predictive power in the ROI were large for surprisal in both paradigms (single words: *d* = 1.62; connected speech: *d* = 2.14) but for entropy only in connected speech (*d* = 1.72) and not in single words (*d* = 0.39).

To test that this effect was not due to the unequal amounts of data in the two experiments, we performed a follow-up analysis with a subset of the continuous-speech data. Stimulus segments 5 and 6 of the continuous-speech experiment together contained 4964 phonemes, comparable to the 4889 phonemes in the single-word experiment. **Figure 4** shows the comparison between experiments when number of phonemes was matched. As expected, this reduction in the amount of data led to a reduction in effect sizes for continuous speech (surprisal: *d* = 1.45; entropy: *d* = 1.36), but it did not change the main result: the ratio between entropy and surprisal was still higher in the continuous-speech data than in single words (continuous speech: *M* = 1.01, *SD* = 0.73; vs. single words: *t_28_* = 3.67, *p* = .001).

**Figure 4.**
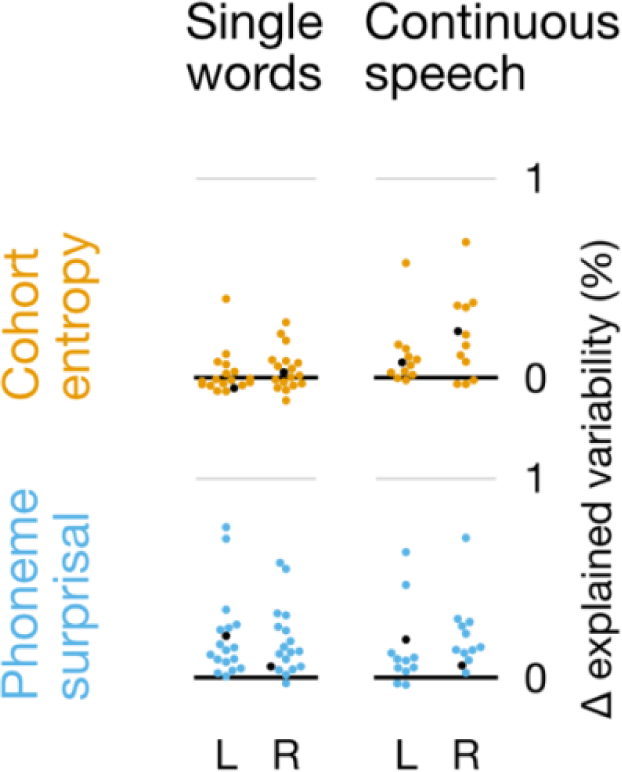
Comparison Between Single Words and Continuous Speech When Number of Phonemes is Matched Note. Matching the number of phonemes between experiments leads to more comparable effect sizes, but does not change the primary conclusions. Details are the same as in **Figure 3B, C**, but using only a subset of the data from the continuous-speech experiment, so that the number of phonemes heard by each participant is matched between the two experiments.

A further concern is that isolated monosyllabic words may be too short to engage higher-level processes. If it is only multisyllabic words that engage processes giving rise to cohort entropy effects, then this could lead to an imbalance between entropy and surprisal effect size in the whole single-word dataset. To address this, we re-analyzed the single-word data with separate entropy and surprisal predictors for mono- and multisyllabic words (the stimuli contained 453 mono- and 547 multisyllabic words). The overall predictive power was reduced, as expected due to the reduced amount of training data, especially for monosyllabic words (which contain fewer phonemes). However, the overall pattern remained the same, with higher predictive power for surprisal than entropy (average in the ROI, monosyllabic: *t_28_* = 2.42, *p* = .027; multisyllabic: *t_28_* = 5.06, *p* < .001).

Finally, we examined the nature of the estimated response functions for phoneme surprisal in the single-word dataset **(Figure 5)**. The analysis of the TRFs was restricted to a mirror-symmetric anatomical region, based on the area in which surprisal significantly improved the model fit in at least one hemisphere. The overall TRF amplitude exhibited two broad peaks, centered on approximately 100 and 350 ms latency (**Figure 5A**). The anatomical distribution of estimated currents in both peaks is consistent with primary sources in the bilateral superior temporal gyrus (**Figure 5B**). In order to visualize the direction of the source currents, we extracted the first principal component of the TRF for each participant and each hemisphere (**Figure 5C, D**). **Figure 5D** shows the average anatomical distribution of the first principal component across subjects. The result in both hemispheres is consistent with a current dipole in auditory cortex, whose average direction is indicated by the arrows in **Figure 5D**. The corresponding time-course, visualized in **Figure 5C**, indicates that the early peak had an upward current direction, while the second peak was dominated by downward current. This time course was further analyzed with mass-univariate *t*-tests, correcting for the time range from 0 to 1000 ms. Even though activity in the early peak did not reach significance in the right hemisphere, the difference between hemispheres, was not significant (*p* = .063, at 70 ms).

## Discussion

This study examined cohort entropy and phoneme surprisal effects in a single-word paradigm using an mTRF analysis, modeling both acoustic and linguistic predictors of neural activity. We found that phoneme surprisal is a significant predictor of neural activity during speech recognition, as have many previous studies (Brodbeck et al., 2018, 2022b; Donhauser & Baillet, 2020; Ettinger et al., 2014; Gagnepain et al., 2012; Gaston & Marantz, 2018; Gillis et al., 2021; Gwilliams et al., 2020; Gwilliams & Marantz, 2015). The spatial distribution of the effect along the superior temporal gyrus is also consistent with previous work. The TRF for phoneme surprisal in our study appears to peak twice, in line with Gwilliams and Marantz (2015), Gaston and Marantz (2018), and Brodbeck et al. (2022b).

In contrast to the robust effect of phoneme surprisal, we did not observe a significant effect of cohort entropy. In a direct comparison to our single-word dataset, we analyzed an existing continuous-speech dataset (Brodbeck et al., 2022b) in the same manner, and found effects of both phoneme surprisal and cohort entropy. The ratio of the predictive power between entropy and surprisal differed significantly between the two experiments. The direct comparison of these two datasets substantiates our generalization about the existing literature, that cohort entropy effects are weak or non-existent in studies that use single words or short phrases, while they are robust in studies that use continuous, naturalistic speech as stimuli.

How could this dissociation between phoneme surprisal and cohort entropy occur? As reviewed in the Introduction, it is frequently assumed that speech input triggers a relatively automatic and uniform process including incremental activation of phoneme, wordform, lexical-syntactic, and conceptual units. However, if the same neural process is engaged for word recognition in single words and in continuous speech, then the neural response should also reflect the same lexical properties. This would predict cohort entropy effects for any task involving word recognition. If anything, prevailing assumptions might lead one to expect that lexical uncertainty would be *lower* when additional context is available (potentially minimizing cohort entropy effects in continuous speech). Importantly, however, cohort entropy depends not only on the number of lexical candidates but the distribution of probability among them, and so should not be systematically impacted in this way even when context is accounted for. To make sense of the dissociation that we observed, with stronger cohort entropy effects in continuous speech, in the following sections we hypothesize that (1) brain responses related to phoneme surprisal and cohort entropy arise from different levels of representation or different sub- processes and (2) their dissociation implies a break in the automatic sequence of processing involved in word recognition.

### Non-automaticity in the Lexical Access Sequence

The pattern of dissociation that we observed could have several different explanations, contingent on the precise neural processes indexed by cohort entropy and phoneme surprisal. In **Figure 6A**, we reproduce our illustration in **Figure 1** of a fully automatic processing sequence in response to each phoneme of speech input. In **Figure 6B-D**, we illustrate alternatives to this sequence that might better represent what occurs incrementally in single-word paradigms that do not elicit cohort entropy effects. It is possible that the decoupled processes do not occur at all in single-word processing; alternatively, they could be engaged sporadically, engaged much later (beyond the 1000 ms window captured by the TRF), or engaged in a less strictly incremental, time-locked manner rather than on a phoneme-by-phoneme basis.

**Figure 5.**
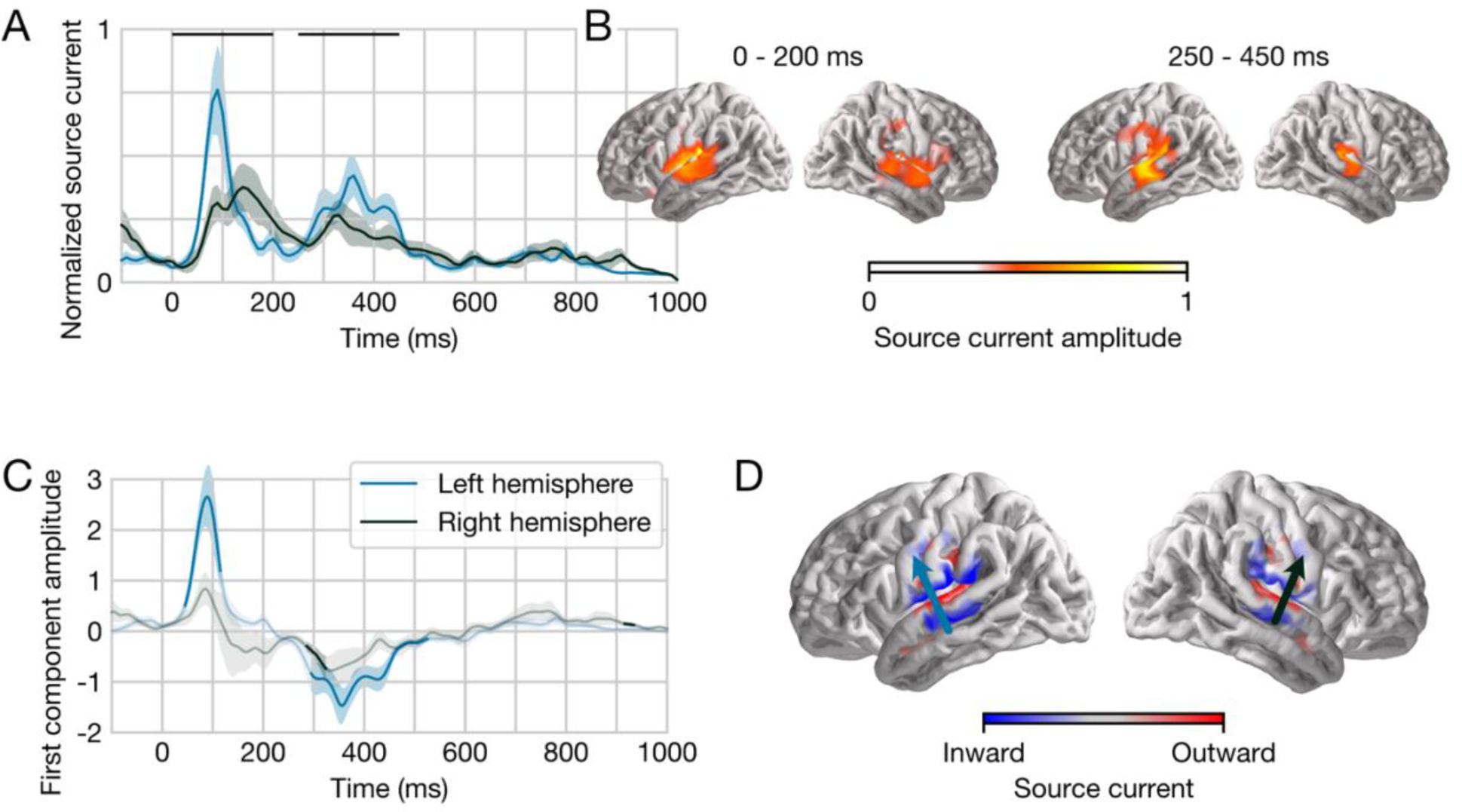
Temporal Response Function (TRF) to Phoneme Surprisal in Isolated Words Note. **(A)** Response amplitude, plotted separately for each hemisphere, summed across all sources in a symmetric region of interest (ROI) defined from significant model improvement due to surprisal in at least one hemisphere. Responses are shown at the normalized scale used for model fitting and with arbitrary units. TRFs exhibit two broad peaks in time. The two black horizontal bars indicate time windows for anatomical plots in panel B. **(B)** Average response amplitude during two peaks in the TRF, suggesting primary sources in the superior temporal gyrus of both hemispheres. Unlike the other plots in this figure, plots in this panel are not constrained to the ROI. **(C)** To visualize current direction over time, the TRF from each subject was decomposed using principal component analysis, separately for the left and the right hemisphere. The plot shows the average time course of the first principal component across subjects. Opaque line segments indicate time ranges in which the respective TRF is significantly different from zero. **(D)** Average anatomical distribution of the first principal component. The color indicates current on the cortical surface, directed into or out of the brain. The average current direction for each hemisphere, indicated by the arrows, is consistent with auditory cortex activity.

**Figure 6.**
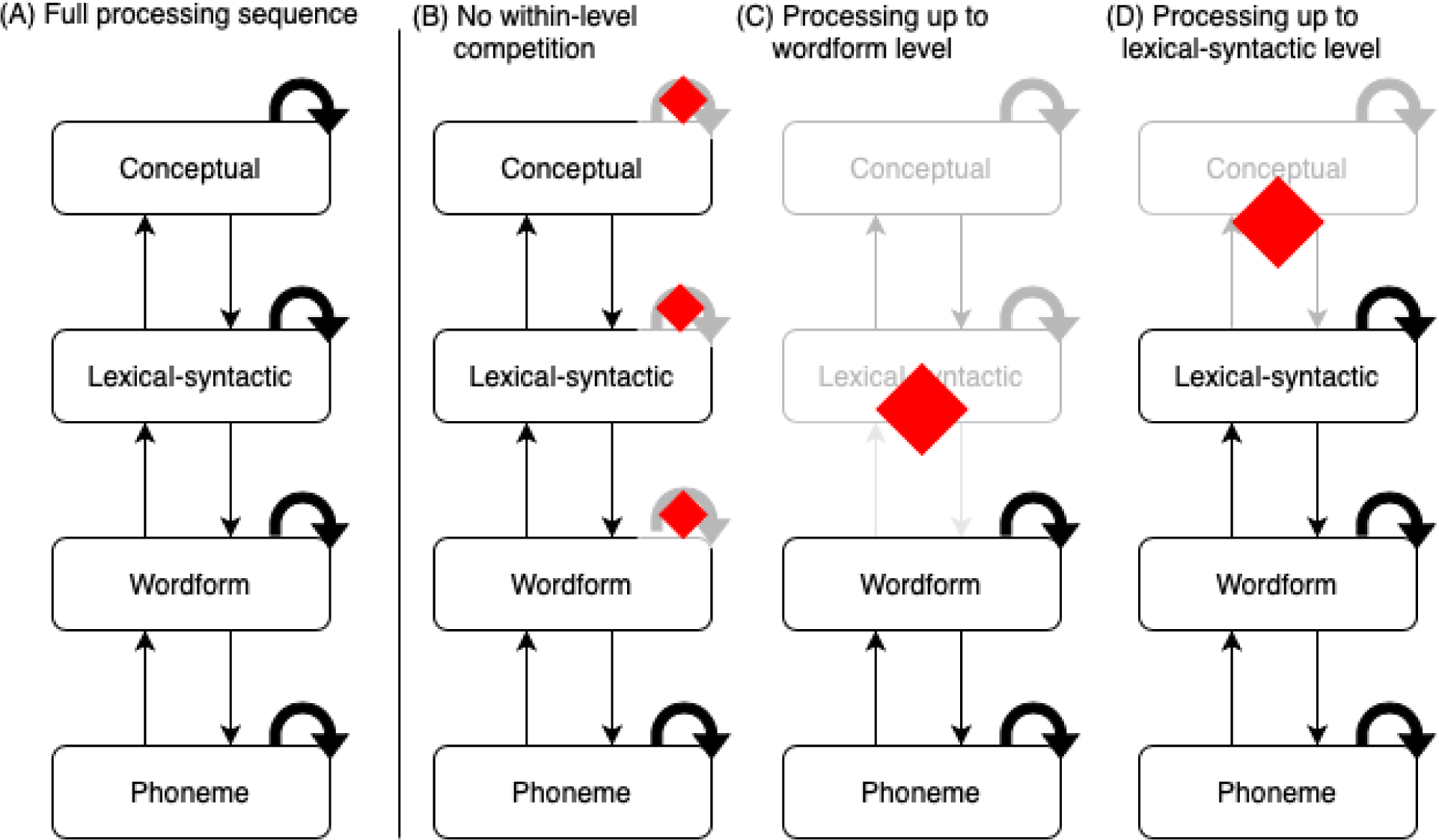
Fully Automatic vs. Alternative Processing Sequences in Response to Each Phoneme of Speech Input Note. **(A)** Fully automatic processing sequence in which both phoneme surprisal and cohort entropy effects arise. **(B)-(D)** Proposed partial processing sequences in which phoneme surprisal but not cohort entropy effects occur. Red diamonds indicate processes or levels of representation that might be delayed or suspended from incremental (phoneme-locked) processing during recognition of single words. As in ***Figure 1***, straight arrows indicate connections between levels of representation. Curved arrows indicate a within-level competition/selection process.

One possible explanation for the dissociation is based on the reasoning that cohort entropy is specifically a measure of the amount of lexical competition occurring (Gagnepain et al., 2012). We can imagine a scenario in which initial activation of multiple lexical candidates is automatic, but the competitive process of winnowing out the weaker candidates is only applied when rapid selection of a single best candidate is particularly helpful or necessary for the task at hand. Accordingly, phoneme surprisal effects might require only *activation* of, e.g., the wordform level of representation, rather than the competition process that occurs within that level (a scenario illustrated by **Figure 6B**, in which within-level competition processes are not occurring above the phoneme level). In contrast, cohort entropy effects would reflect the competitive selection process that allows a single best candidate to be identified as early as possible, and this process might only be engaged when faced with the time pressure of processing connected speech.

Another possibility is that phoneme surprisal and cohort entropy effects reflect different levels of representation which are not all automatically accessed to the same degree. Access to ’lower’ levels of representation like phoneme or wordform representations might be more automatic, whereas access to ‘higher’ levels of representation like lexical-syntactic or conceptual units might be more dependent on context and task demands. For instance, surprisal effects might require only wordform-level activation, while cohort entropy effects might require lexical-syntactic activation or higher. Similarly, phoneme surprisal effects could implicate up to lexical-syntactic activation while cohort entropy effects require conceptual activation or higher. These two possibilities are illustrated in **Figure 6C** and **Figure 6D**, respectively. Consistent with such an explanation, semantic priming from partial wordforms seems to be more reliable in connected speech (Zwitserlood, 1989) than in a single-word lexical-decision paradigm, where form-based priming predominates (Gaskell & Marslen-Wilson, 2002). Even within a single-word paradigm, Bentin et al. (1993) argue that the extent to which a task requires semantic processing can influence the degree of semantic priming that occurs, as indexed by the N400 response.

Though less likely, we can acknowledge two alternative explanations in which phoneme surprisal effects do not reflect the activation of wordform representations, while cohort entropy effects do. We consider these explanations less likely because they would imply an absence of incremental wordform-level processing in the single-word tasks, despite behavioral evidence to the contrary. One possibility is that apparent phoneme surprisal effects arise due to prelexical phonotactic processing, involving representations sensitive to the probability of phoneme sequences in the language independent of wordform representations. The second possibility arises from the proposal of Norris & McQueen (2008) that ‘off-line’ perceptual learning could lead to wordform frequency effects on phoneme probability without concurrent wordform activation causing online top-down effects. In either of these scenarios, a phoneme surprisal effect does not necessarily imply wordform activation; cohort entropy could reflect anything at the wordform level or above.

It also remains possible that correlations between neural activity and cohort entropy are not driven by lexical competition or uncertainty per se but by a secondary process that is sensitive to lexical competition or uncertainty. If that process is only engaged by continuous speech, cohort entropy effects would also appear to be modulated, despite lexical processing in either case. A clear contender for such a secondary process emerges from outside the domain of language. Neural effects of entropy have been hypothesized to reflect heightened sensitivity, via increased post-synaptic gain, to bottom-up sensory inputs in high-uncertainty contexts (Donhauser & Baillet, 2020; Feldman & Friston, 2010; Friston, 2005; Strange et al., 2005). To maintain this hypothesis while integrating our findings, it must be admitted that this top-down modulation is not obligatory: the effect of surprisal in our study provides evidence that linguistic representations above the phoneme were engaged, and yet we did not see the effect of entropy, associated with the hypothesized sensitization to the bottom-up input. Our results do not directly speak to whether the entropy effects arise due to sensitization to the input, or due to neural activity at a different level of processing. However, in either case, the results suggest that there is a neural process that leads to this modulation of brain activity in continuous speech, but not in single words.

### Single Words vs. Continuous Speech

What are the differences between single-word paradigms and continuous speech that would make any of the distinctions described in the previous section possible? First, the reliable presence of pauses between words in single-word studies may constitute a key change in task demands, by leaving sufficient time for full lexical access to occur after wordform offset and before the next wordform begins and thus reducing the necessity for incremental processing.

Early competitive selection might be unnecessary, and/or access to higher-level syntactic and conceptual units could be deferred until the pause makes the auditory wordform uniquely identifiable. Among the single-word studies we have reviewed, the pause detection (Gagnepain et al., 2012) and nonword detection (Kocagoncu et al., 2017) tasks incorporate lengthy inter-stimulus intervals averaging 2000 ms, and the lexical decision studies (Brennan et al., 2014; Ettinger et al., 2014; Gwilliams & Marantz, 2015; Lewis & Poeppel, 2014) wait for a participant response after each word. Our study used a shorter but still considerable inter-stimulus interval of 267 ms with only occasional semantic relatedness probes, and also did not find a cohort entropy effect.

Second, the syntactic and semantic structure in continuous speech provides another motivation for incremental processing: rapid access to lexical and conceptual content for the current word provides information that might aid recognition of the subsequent word. This rationale for rapid processing is absent in single-word paradigms that lack structure. Even beyond not requiring speed in lexical or conceptual access, the tasks employed in single-word paradigms may in some cases not require lexical or conceptual access at all. For instance, our task involved semantic relatedness judgements with written probes. It is conceivable that this task might be solved successfully by temporarily ‘buffering’ the input from each word as a form-based representation, and only accessing conceptual representations if a probe occurs, rather than accessing lexical and conceptual representations for every stimulus. By contrast, the speed of continuous speech, its many between-word dependencies, and the imperative to build sentence-level and message-level interpretations could be what drive competition or incremental higher-level activation (and therefore cohort entropy effects) in naturalistic paradigms.

We might expect that cohort entropy effects could be observed for single words if a task were designed such that earlier identification of the word is encouraged and incremental higher-level activation becomes more advantageous, whether via the elimination of pauses or the addition of some higher-level structure. Likewise, pauses could be added to continuous speech. The three-word phrases used by Gaston and Marantz (2018) (e.g., “to chew gum,” “the shredder broke”) are an interesting test of these hypotheses, as they lack within-phrase pauses and have syntactic and semantic structure. Nevertheless, Gaston and Marantz did not find cohort entropy effects when their cohort entropy variables were evaluated in the same model as phoneme surprisal. This suggests that only longer sequences of continuous speech elicit cohort entropy effects, and therefore that a buffering process may play a mediating role here. Speech rate (almost certainly higher in studies using continuous speech) may also be relevant.

Another line of investigation for understanding what drives neural cohort entropy effects might involve the contrast between monomorphemic and multimorphemic words. The two types of stimuli can be closely matched in length (itself another factor to consider), but only multimorphemic words can be viewed as structured sequences of units of meaning of the kind that might encourage more incremental processing. The inclusion of multimorphemic words in a single-word study could thus motivate earlier selection and higher-level activation so that initial morphemes can be recognized in time to begin processing any potential subsequent morphemes. In **Table 1**, we noted that all single-word studies that do not include multimorphemic words also do not report cohort entropy effects (Brennan et al., 2014; Gagnepain et al., 2012; Lewis & Poeppel, 2014). This is true of our study as well. Among the single-word studies that do report cohort entropy effects, albeit without controlling for phoneme surprisal, Ettinger et al. (2014) include multimorphemic words and Kocagoncu et al. (2017) do not indicate whether multimorphemic words are included in their stimuli or not. This factor deserves further investigation.

For any potentially relevant stimulus properties that might distinguish isolated words and continuous speech and that can be conceptualized as continuous variables, attempting to demonstrate co-variation between this stimulus property and the size of the entropy effect *within* a dataset would be a particularly effective means of narrowing the current hypothesis space.

### Implications

If auditory word recognition in most single-word studies proceeds in the manner we have proposed, with candidate selection or higher-than-wordform-level processing delayed or suspended entirely, there are two major implications. The first is that the cascading, incremental lexical access process is not automatic but rather is motivated by time pressure and modulable with the extent of that time pressure. The second is that auditory word recognition in many single-word studies may differ fundamentally from the process most researchers assume they are studying (that is, speech recognition in natural contexts). This would invite re-interpretation of existing neural and behavioral data and would motivate increased use of more naturalistic designs in future work, or identification of changes to single-word paradigms that would drive cohort entropy effects so that these paradigms can be used with more confidence that they are representative of the processing of natural, connected speech. We note that some of the strongest behavioral evidence for incremental lexical processing comes from eye-tracking in the visual world paradigm, as this method provides a continuous measure of lexical activation over the course of speech input. Most visual world studies, however, present target lexical stimuli within sentences, and even with single-word stimuli the task (find a visual referent) may encourage rapid incremental access. Reconciliation between apparent cohort activation effects in the visual world paradigm and neural cohort entropy effects is likely to be a productive step forward, integrating paradigms that investigate the status of individual items in order to make inferences about the set of lexical candidates and paradigms that investigate properties of the set itself.

## Conclusion

Our goal in this study was to evaluate whether auditory word recognition should be assumed to proceed in the same way for isolated single words and natural connected speech. We also intended to establish a better understanding of what drives phoneme surprisal and cohort entropy effects during speech recognition, while modeling the stimulus as thoroughly as current methods allow. We directly compared single-word and continuous-speech data from MEG and demonstrated that the paradigms differ in a way that is consistent with patterns in the existing literature, where phoneme surprisal effects are robust and cohort entropy effects occur sporadically. We found that the ratio between the predictive power of cohort entropy and phoneme surprisal is significantly higher (closer to one) for continuous speech as compared to single words; indeed, we do not observe a cohort entropy effect for single words at all. We proposed that this is because phoneme surprisal effects arise from the activation of a lower-level (e.g., wordform) representation of the speech input while cohort entropy effects arise from a competition process or activation of a higher-level (e.g., lexical-syntactic) representation whose engagement is delayed or does not occur in single-word paradigms. This dissociation suggests (1) that the full sequence of steps involved in auditory word recognition does not proceed automatically and (2) that the extent to which higher levels of representation and/or lexical competition processes are engaged depends on the nature of the stimulus or experimental task. Finally, this study has also helped validate multivariate temporal response function analysis as a promising method for future work in single-word paradigms.

